# Targeting APEX2 to the mRNA encoding fatty acid synthase β in yeast identifies proteins that bind and control its translational efficiency in the cell cycle

**DOI:** 10.1101/2023.05.02.539120

**Authors:** Heidi M. Blank, Wendell P. Griffith, Michael Polymenis

**Affiliations:** Department of Biochemistry and Biophysics, Texas A&M University, College Station, TX 77843, USA; Department of Chemistry, The University of Texas at San Antonio, San Antonio, TX 78249, USA

**Keywords:** Cas13, Tdh3p, FAS1, proximity labeling

## Abstract

Profiling the repertoire of proteins associated with a given mRNA during the cell cycle is unstudied. Furthermore, it is much easier to ask and answer what mRNAs a specific protein might bind to than the other way around. Here, we implemented an RNA-centric proximity labeling technology at different points in the cell cycle in highly synchronous yeast cultures. To understand how the translation of *FAS1*, encoding fatty acid synthase, peaks late in the cell cycle, we identified proteins that bind the *FAS1* transcript in a cell cycle-dependent manner. We used dCas13d-APEX2 fusions to target *FAS1* and label nearby proteins, which were then identified by mass spectrometry. The glycolytic enzyme Tdh3p, a known RNA-binding protein, bound the *FAS1* mRNA, and it was necessary for the increased Fas1p expression late in the cell cycle. Lastly, cells lacking Tdh3p had altered size homeostasis, consistent with delayed G1/S transition and exit from mitosis. These results point to unexpected connections between major metabolic pathways. They also underscore the role of mRNA-protein interactions for gene expression during cell division.

## INTRODUCTION

Proteins that bind to each mRNA could influence multiple steps in gene expression, impacting the mRNA’s processing, stability, or interaction with ribosomes and translation. The repertoire of protein-mRNA interactions has been traditionally defined from protein-centric methods, tagging a given mRNA binding protein (mRBP), and answering what mRNAs bind to the mRBP (Hogan *et al*, 2008). The converse, mRNA-centric approach to identify what proteins a specific mRNA binds is challenging because it requires tagging the mRNA of interest. Recently, new technologies, including engineered CRISPR-Cas systems, have been implemented to target particular mRNAs (Han *et al*, 2020; Li *et al*, 2021; Abudayyeh *et al*, 2017). When combined with ascorbate peroxidase (APEX)-based or similar proximity-labeling tools, proteins interacting with the RNA of interest can be identified (Li *et al*, 2021; Han *et al*, 2020).

There are several unexplored contexts where identifying mRNA-mRBP interactions could offer significant biological insight. For example, in recent years, ribosome profiling experiments have identified mRNAs that are translated with different efficiency during cell division in bacterial (Schrader *et al*, 2016), human (Tanenbaum *et al*, 2015; Stumpf *et al*, 2013) or yeast cells (Blank *et al*, 2017b; Maitra *et al*, 2020). A key question is how translational control can be imposed when protein synthesis rates remain unchanged as cells progress in the cell cycle (Tanenbaum *et al*, 2015; Elliott & McLaughlin, 1978; Stonyte *et al*, 2018). Changes in ribosome abundance resulting, for example, from nutrient changes, impose straightforward translational control on specific mRNAs (Mills & Green, 2017). But the ribosome content does not change in the cell cycle (Blank *et al*, 2020; Elliott *et al*, 1979). On the other hand, specific mRBP-mRNA interactions could establish translational control during cell division.

Several mRBPs have altered levels in the cell cycle, their loss-of-function mutations lead to cell cycle phenotypes, or they are targeted by the cyclin-dependent kinase (Cdk) that drives cell cycle transitions (Polymenis, 2022a). However, there are few examples of mRBP-mRNA interactions with known roles during cell division. The best case in budding yeast is Whi3p, which binds the G1 cyclin *CLN3* mRNA. The Whi3p-*CLN3* interactions destabilize *CLN3* and also repress its translation (Cai & Futcher, 2013). In mammals, the DENR-MCT1 heterodimer is phosphorylated by mitotic Cdk/cyclin complexes, enabling it to derepress translation of specific mRNAs needed for the proper execution of mitosis (Clemm von Hohenberg *et al*, 2022). These examples notwithstanding, there is little additional information on mRBP-mRNA interactions significant for cell division. To fill this gap in knowledge, it is necessary to examine dynamic mRBP-mRNA interactions in highly synchronous cells as they progress in the cell cycle.

Past work probing RNA-protein interactions revealed several metabolic enzymes moonlighting as RNA-binding proteins (Beckmann *et al*, 2015; Hentze *et al*, 2018). For example, the glycolytic enzyme GAPDH, which converts glyceraldehyde-3-phosphate and NAD^+^ to 1,3-bisphosphoglycerate and NADH, binds to multiple RNAs in mammalian (Dollenmaier & Weitz, 2003; Ryazanov, 1985; Singh & Green, 1993; Rodríguez-Pascual *et al*, 2008; Bonafé *et al*, 2005; Castello *et al*, 2016) and yeast (Mitchell *et al*, 2013; Hogan *et al*, 2008; Matia-González *et al*, 2015) cells. These enzyme-RNA interactions may influence not only the target RNA but also the enzyme’s catalytic activity (e.g., by blocking access to its metabolite substrates), leading to the formulation of the RNA-enzyme-metabolite (REM) hypothesis of metabolic and gene expression control (Hentze & Preiss, 2010). Nonetheless, in most cases, the physiological function of the RNA-binding activity of metabolic enzymes and whether they regulate their target mRNAs remain unknown (Castello *et al*, 2015).

Here, we describe the application of RNA-centric proximity labeling in yeast and the first cell cycle-dependent interrogation of mRBP-mRNA interactions in any system. We had previously reported the identification of mRNAs with altered translational efficiency in the cell cycle (Blank *et al*, 2017b). Among these mRNAs was *FAS1*, encoding the β subunit of fatty acid synthase. The translational efficiency of *FAS1* and the levels of the Fas1p protein peak late in the cell cycle, providing lipid resources needed for mitosis (Blank *et al*, 2017b; Maitra *et al*, 2022; Blank *et al*, 2017a). We implemented dCas13d-APEX2 mediated proximity labeling to identify proteins interacting with the *FAS1* mRNA in a cell cycle-dependent manner. Our results suggest that Tdh3p, a yeast GAPDH isoform, binds *FAS1* and it is necessary to promote Fas1p synthesis late in the cell cycle. These results reveal unexpected gene expression control layers during cell division. Furthermore, they point to possible connections between enzymes of major metabolic pathways, such as fatty acid synthesis (Fas1p) and glycolysis (Tdh3p). Lastly, the approaches we used can apply to other systems.

## RESULTS AND DISCUSSION

### Generating active dCas13d-APEX2 targeting the *FAS1* mRNA in yeast

To bring APEX2 to the *FAS1* transcript, we decided to deploy in yeast the CRISPR-Cas13d targeting approach reported recently for mammalian cells (Han *et al*, 2020). Although dCas13d (encoding a catalytically dead, guide-RNA directed ribonuclease) targets exclusively RNA (Zhang *et al*, 2018), for proximity labeling applications, the interaction is stabilized by adding a double-stranded RNA binding domain (dsRBD) (Han *et al*, 2020). To drive the expression of dCas13d-dsRBD-APEX2 in yeast, we placed this construct (C-terminally tagged with the V5 epitope for protein surveillance purposes) under the control of a strong promoter (Figure 1A). In the same integrative plasmid, we also placed in a bicistronic arrangement the necessary sequences for the expression of a guide-RNA (Figure 1A; see Materials and Methods). We chose three predicted guide-RNA sequences for Cas13 systems (Wessels *et al*, 2020), targeting the long *FAS1* transcript (∼6.5-7 kb) at the positions shown in Figure 1B (see also Materials and Methods). Each of the three bicistronic constructs, carrying both the dCas13d-dsRBD-APEX2-V5 and one of the guide-RNA cistrons, was then integrated at the *URA3* locus, generating three yeast strains (*FAS1-1*, *FAS1-2*, and *FAS1-3*) that were used in our subsequent proximity labeling experiments.

**Figure 1.**
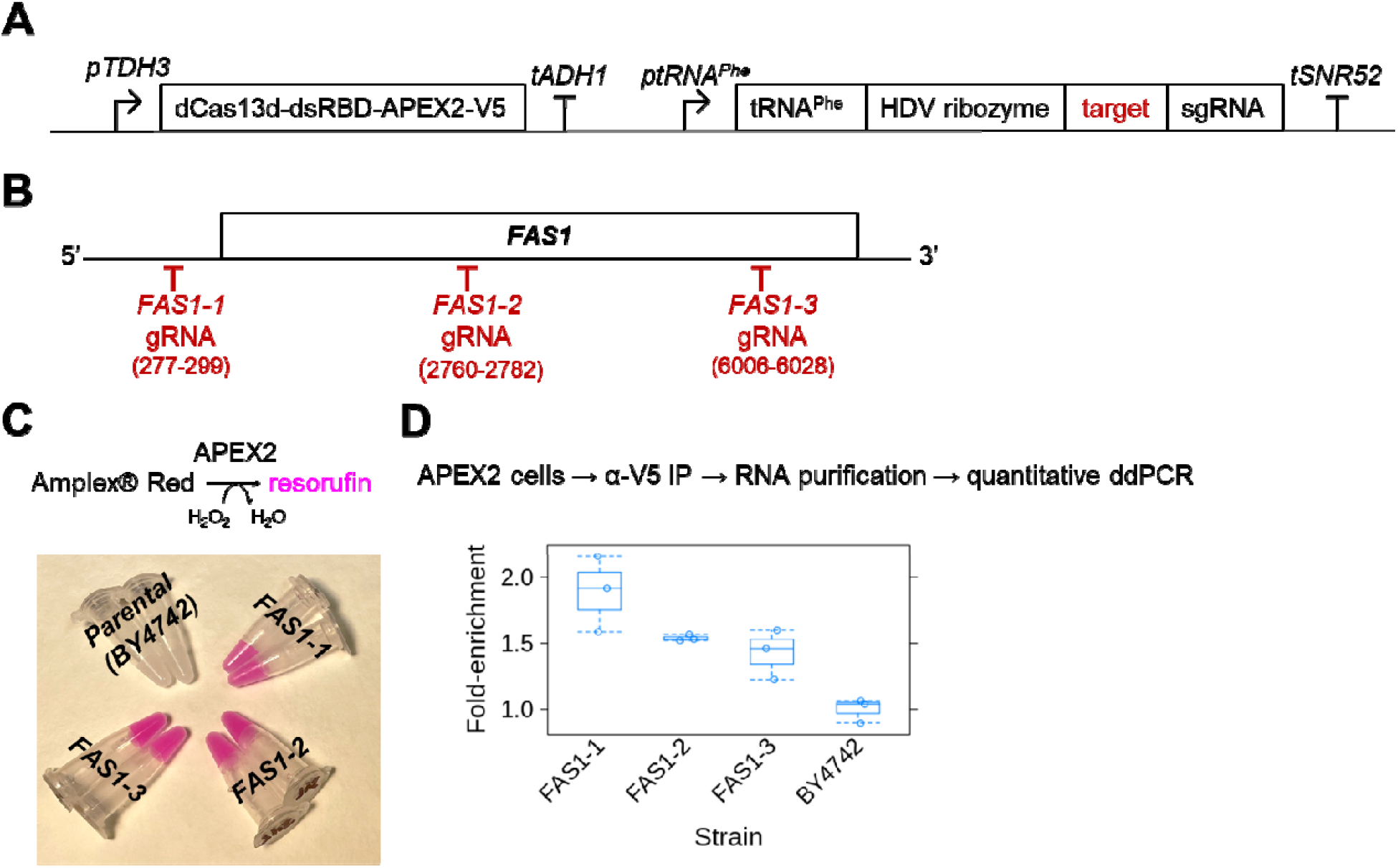
Engineered yeast cells express active dCas13d-APEX2 targeting the *FAS1* mRNA. A, Diagram of the engineered bicistronic locus introduced into yeast cells to express dCas13d-APEX2 and gRNAs targeting *FAS1* (see Materials and Methods). B, Schematic of the targeted positions on the *FAS1* mRNA. C, Cells of the indicated genotype, carrying different gRNAs targeting *FAS1*, express active APEX2 based on the conversion of Amplex^Ⓡ^ Red to resorufin. The cells were processed as described in Materials and Methods. D, Yeast cells expressing dCas13d-dsRBD-APEX2-V5, see (A), target it preferentially to the *FAS1* mRNA. Digital droplet PCR (ddPCR; see Materials and Methods) was used to measure the levels of *FAS1* immunoprecipitated by dCas13d-dsRBD-APEX2-V5 shown on the y-axis in the strains shown on the x-axis. The values used to generate the graphs are in File S1/Sheet1.

To test if ascorbate peroxidase activity is present in the yeast cells, we exposed them to hydrogen peroxide and Amplex^Ⓡ^ Red, which in an APEX2-catalyzed reaction is oxidized to the fluorescent product resorufin (Dębski *et al*, 2016). Cells from all three strains expressing dCas13d-dsRBD-APEX2-V5 along with a *FAS1* guide-RNA became highly fluorescent compared to the parental strain that does not express APEX2 (Figure 1C). Furthermore, using the V5 epitope, we immunoprecipitated dCas13d-dsRBD-APEX2-V5 and asked if the *FAS1* mRNA was associated with it, as measured by digital droplet PCR (ddPCR; see Materials and Methods). We found a moderate (∼1.5-2-fold) but significant (p<0.0001, based on the robust bootstrap ANOVA; see Materials and Methods) enrichment of *FAS1* levels in the immunoprecipitated samples compared to the input levels in the cell extracts (Figure 1D). We note that in the published example of dCas13d-dsRBD-APEX2-V5 targeted to the human telomerase RNA, the reported enrichment was 3-4-fold (Han *et al*, 2020). These results suggest that dCas13d-dsRBD-APEX2-V5 was active in yeast cells and targeted the *FAS1* transcript.

### Establishing labeling conditions in cells expressing dCas13d-APEX2 fusions

Our next objective was to establish the conditions necessary to observe labeling, reported by the appearance of biotinylated proteins. APEX labeling has not been very successful in yeast because yeast cells are impermeable to biotin-conjugated phenol, which needs to be oxidized by the APEX peroxidase into its reactive, free radical form before it can form covalent bonds with electron-rich groups, such as those found on the side chains of amino acids. It has been reported that weakening the cell wall with zymolyase (Hwang & Espenshade, 2016) or osmotic shock with freeze-thaw cycles (Singer-Krüger *et al*, 2020) improves APEX-mediated labeling. An alternative strategy relies on the cellular uptake of a chemical probe that affixes to targets via APEX labeling. The chemical group is then derivatized with click chemistry in vitro, in cell extracts, to attach biotin to the labeled targets (Li *et al*, 2020). We tried all the above procedures, but with limited success. Osmotic shock with freeze-thaw cycles did improve the observed labeling, based on the appearance of biotinylated proteins on immunoblots (Figure 2; compare the first lane revealing the endogenous biotinylated yeast proteins to the second lane revealing the increased APEX-mediated labeling). To achieve more efficient labeling, in addition to osmotic shock, we relied on a previously described approach, employing digitonin permeabilization, to measure glycolysis in situ (Cordeiro & Freire, 1995). As we detail in Materials and Methods, permeabilization of cells with digitonin (used at 0.01%) resulted in strong labeling and appearance of biotinylated proteins (Figure 2; see last five lanes). These results argue that we had in place the necessary tools and experimental conditions to identify the proteins interacting with the *FAS1* transcript.

**Figure 2.**
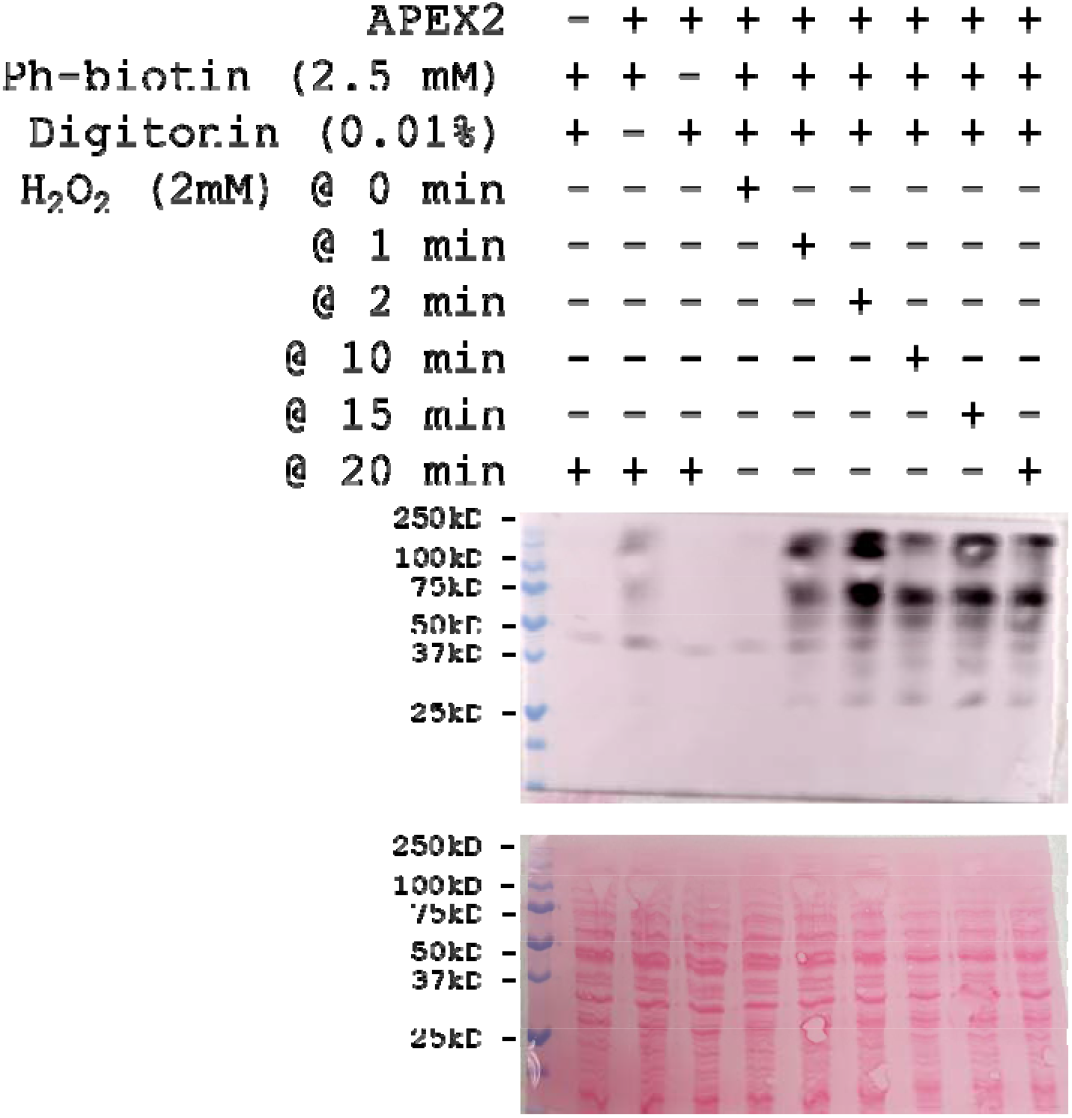
Biotin labeling conditions in cells expressing dCas13d-APEX2 fusions. The immunoblot displays the signal from biotinylated proteins in cells treated in each condition shown on top. The first lane is from extracts prepared using strain BY4742, and the rest is from extracts using strain *FAS1-1.* The blot at the bottom is the one shown above before it was processed for immunodetection, stained with Ponceau S to reveal total protein loading.

### Proximity labeling of proteins targeting *FAS1* in the cell cycle

We relied on centrifugal elutriation to collect synchronous cells because it is a selection method that is less disruptive of the normal coordination between cell growth and division than arrest-and-release methods (Aramayo & Polymenis, 2017; Polymenis, 2022b). To overcome the low yield associated with elutriation, we generated pools of cells collected at the same cell size, as we have done in the past (Blank *et al*, 2017b; Maitra *et al*, 2020). Because yeast daughter cells actively monitor their size to adjust progression in the cell cycle (a.k.a. ‘sizer’ behavior) (Di Talia *et al*, 2007), their position in the cell cycle is reflected by how big they are. From a total of 54 elutriated cultures (Figure 3A), for each of the three engineered strains (FAS1-1, FAS1-2, FAS1-3), we generated three pools of small, unbudded, G1 cells and three pools of large, budded, non-G1 cells (Figure 3B). Each pool consisted of ∼1E+09 cells, and it was processed for APEX proximity labeling as described in Materials and Methods.

**Figure 3.**
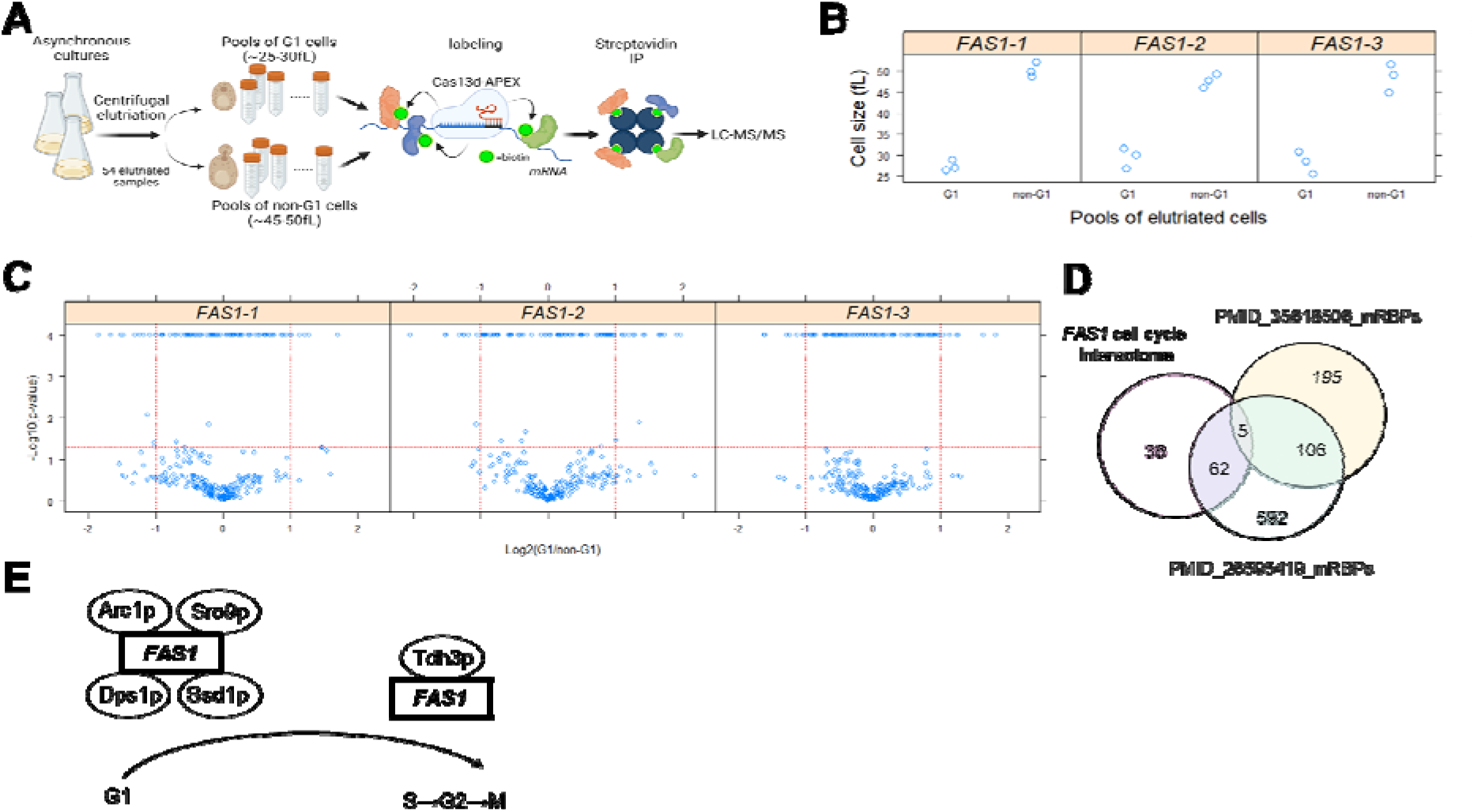
Proximity labeling of proteins targeting *FAS1* in the cell cycle. A, Schematic overview of our experimental approach. B, The cell size (y-axis) of the pools of cells we isolated from each strain is shown for the G1 and non-G1 cells (x-axis). The values used to generate the graphs are in File S1/Sheet2. C, Volcano plots depicting the proteins identified by mass spectrometry in the indicated strain (shown above each panel) whose levels changed significantly in G1 vs. non-G1 cells, based on the magnitude of the difference (x-axis; Log2-fold change) and statistical significance (y-axis), indicated by the red lines. The analytical and statistical approaches are described in Materials and Methods. The values used to generate the graphs are in File S1/Sheet3. Note that the lowest calculated p-values from the robust ANOVA were at the 0.0001 level. D, Venn diagram of the proteins we identified to interact with *FAS1* in a cell cycle-dependent manner (left set) against two reference sets (PMID_35618506, right; and PMID_26595419, middle). The values used to generate the graph are in File S1/Sheet4. E, Schematic summary of the mRBPs that bind the *FAS1* transcript in G1 or non-G1 phases.

Biotinylated proteins were immunoprecipitated with streptavidin magnetic beads, digested with trypsin, and subjected to mass spectrometry for protein identification (see Materials and Methods). We identified 937 proteins in the immunoprecipitated samples, including naturally biotinylated yeast proteins, such as Acc1p (Al-Feel *et al*, 1992; Schneiter *et al*, 1999). As a proxy for the relative abundance of the proteins we identified, we used their exponentially modified protein abundance index (emPAI) scores (Ishihama *et al*, 2005). For each of the three strains, we then used robust bootstrap ANOVA to identify proteins whose abundance changed significantly in G1 vs. non-G1 cells (p<0.05 and fold-change≥2; see Figure 3C and File S1/Sheet3). The levels of 52 proteins changed significantly in the immunoprecipitated samples in G1 vs. non-G1 cells (Figure 3D, right set). Another 51 proteins were found exclusively in G1 or non-G1 cells, but not both (Figure 3D, left set), raising the total to 103 putative hits (File S1/Sheet4).

We note that 67 of the 103 proteins we identified were previously included in a list of 765 yeast proteins that bound RNAs in vivo (Matia-González *et al*, 2015), potentially attesting to the power of our approach. However, that reference compendium (PMID:26595419) was rather expansive, including, for example, ribosomal proteins and known DNA helicases (Matia-González *et al*, 2015). Furthermore, since *FAS1* is highly abundant, one could envision spurious interactions among these 103 proteins. Hence, to prioritize our hits for follow-up studies, we also looked at a smaller reference set of 306 yeast mRBPs (PMID:35618506; (Polymenis, 2022a)) based on earlier in vivo RNA interactome studies (Hogan *et al*, 2008; Mitchell *et al*, 2013). Only five of the 103 proteins we identified here to interact in a cell-cycle-dependent manner with *FAS1* were in both reference mRBP sets (Figure 3D). These five proteins were Arc1p, Dps1p, Sro9p, Ssd1p, and Tdh3p.

Arc1p is involved in tRNA delivery and binds tRNAs and methionyl- and glutamyl-tRNA synthetases (Simos *et al*, 1996), while Dps1p is an aspartyl-tRNA synthetase (Sellami *et al*, 1985). Sro9p has a La-motif involved in RNA binding and it is associated with ribosomes (Sobel & Wolin, 1999). Ssd1p binds and represses mRNAs, especially ones involved in cell wall biosynthesis (Bayne *et al*, 2022; Kaeberlein & Guarente, 2002). Tdh3p is one of three glyceraldehyde-3-phosphate isoforms in yeast (McAlister & Holland, 1985). Regarding their binding to the *FAS1* mRNA, the cell cycle-specific enrichment of all five proteins was highly significant (p<0.0001, based on the robust bootstrap ANOVA). Interestingly, except for Sro9p, identified from strain FAS1-1 expressing a guide-RNA from the 5’-UTR of the *FAS1* transcript, all others were identified from strain FAS1-2, which expresses a guide-RNA from the middle of the *FAS1* transcript (Figure 1B). Lastly, while Tdh3p bound the *FAS1* mRNA preferentially late in the cell cycle, all other proteins bound *FAS1* in the G1 phase (Figure 3E).

### Tdh3p binds *FAS1*, and it is required for cell cycle-dependent changes in Fas1p levels

To follow up on the findings from our proximity labeling experiments, we performed the reciprocal, protein-centric experiments to test the mRBP interactions with the *FAS1* transcript. We used strains carrying TAP-tagged alleles of the corresponding genes, immunoprecipitated the epitope-tagged proteins, and asked if *FAS1* levels were enriched in the immunoprecipitates (see Materials and Methods). Except for *DPS1-TAP*, all other strains were commercially available (see Reagent Table), expressing TAP-tagged proteins (Arc1p, Sro9p, Ssd1p, and Tdh3p) of the expected size (not shown). Attempts to generate a *DPS1-TAP* strain were unsuccessful, so we proceeded with the rest. We found that *FAS1* levels were not enriched with the immunoprecipitated Arc1p-TAP, Sro9p-TAP, or Ssd1p-TAP proteins, but *FAS1* was significantly associated with Tdh3p-TAP (Figure 4A; p<0.0001, based on the robust bootstrap ANOVA). These results do not necessarily exclude the possibility that Arc1p, Sro9p, or Ssd1p interact with *FAS1* in cells. These interactions may be transient and missed in immunoprecipitation experiments but detected by APEX-mediated proximity labeling. We also note that previously reported RNA interactomes typically involve a UV crosslinking step (Hentze *et al*, 2018; Mitchell *et al*, 2013; Matia-González *et al*, 2015), to capture weak RNA-protein interaction. We did not use UV crosslinking in our experiments. Nonetheless, since the Tdh3p interactions with *FAS1* were evident in both approaches we used (RNA proximity labeling and protein immunoprecipitations), we focused on Tdh3p’s role in mediating the cell cycle-dependent changes in the abundance of Fas1p.

**Figure 4.**
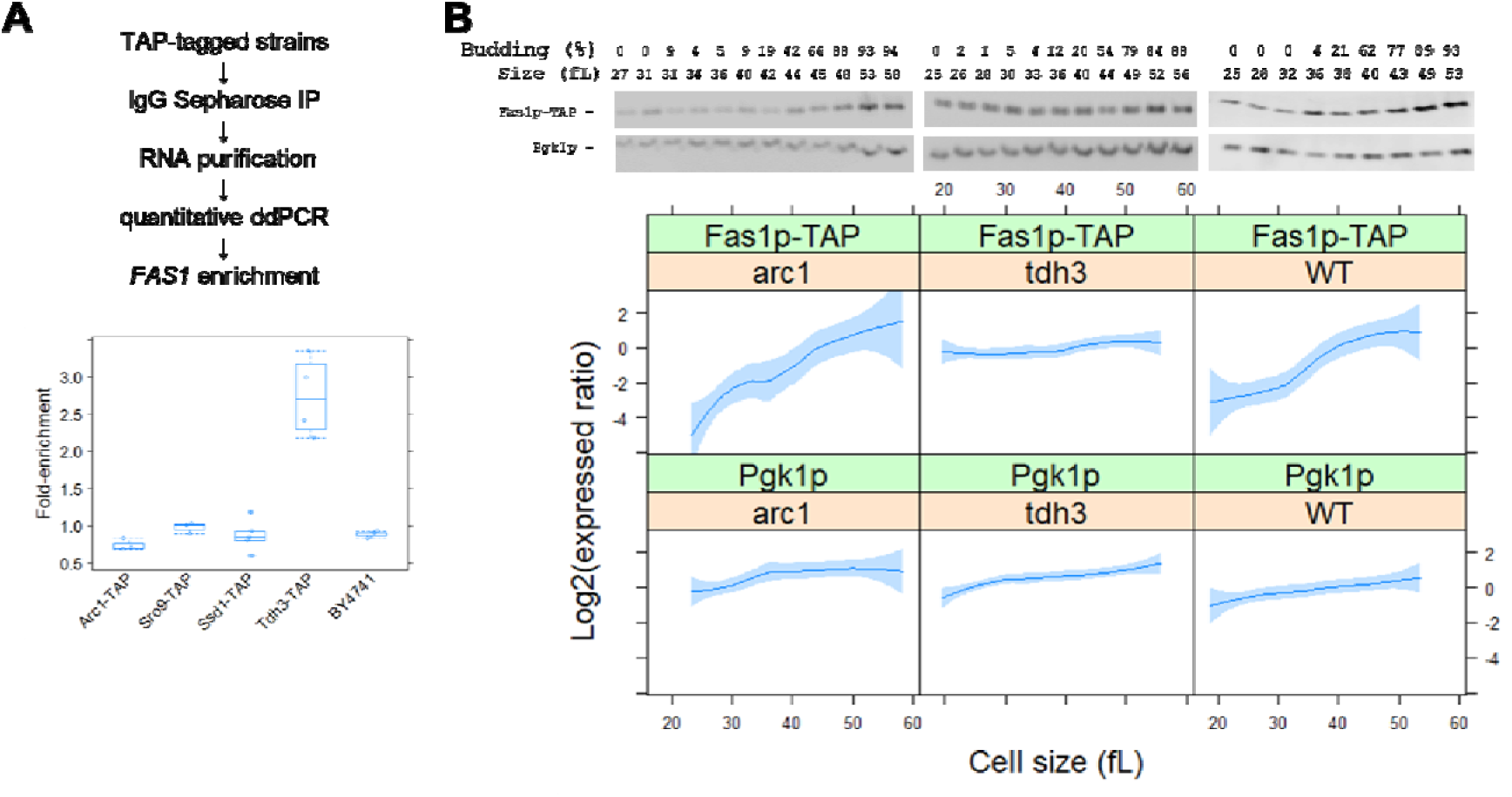
Tdh3p binds *FAS1*, and it is required for cell cycle-dependent changes in Fas1p levels. A, Yeast cells expressing the corresponding TAP-tagged alleles were used to immunoprecipitate the indicated TAP-tagged proteins. The levels of the associated *FAS1* mRNA in the immunoprecipitates (measured as in Figure 1D; see Materials and Methods) are shown on the y-axis in the strains shown on the x-axis. The values used to generate the graphs are in File S1/Sheet5. B, The abundance of TAP-tagged proteins was monitored in strains of the indicated genotype, as described in Materials and Methods. Samples were collected by elutriation in a rich, undefined medium (YPD) and allowed to progress synchronously in the cell cycle. ExperimentlJmatched loading controls (measuring Pgk1p abundance) were also quantified and shown in parallel. (Top), representative immunoblots, along with the percentage of budded cells (% budded) and the cell size (in fL) for each sample. (Bottom), from at least three independent experiments in each case, the TAP and Pgk1p signal intensities were quantified as described in Materials and Methods. The Log2(expressed ratios) values are on the y-axis, and cell size values are on the x-axis. Loess curves and the standard errors at a 0.95 level are shown. All the immunoblots for this figure are in File S3, while the values used to generate the graphs are in File S1/Sheet6.

For Fas1p surveillance in the cell cycle, we used cells carrying a *FAS1-TAP* allele expressed from its native chromosomal locus and introduced a *TDH3* or *ARC1* deletion (the latter used as an additional control in this experiment because we saw no binding of Arc1p to *FAS1* in the immunoprecipitations; see Figure 4A). We then isolated by centrifugal elutriation early G1 cells. We measured their size and budding as cells progressed in the cell cycle and collected samples for immunoblotting at regular intervals. While Fas1p levels increased markedly late in the cell cycle in wild-type and *arc1*Δ cells, they remained constant in *tdh3*Δ cells (Figure 4B). We conclude that Tdh3p is necessary for the cell cycle-dependent changes in Fas1p levels, arguing for a physiological role for the Tdh3p-*FAS1* interactions.

The above experiments allowed us also to evaluate cell cycle kinetics in cells lacking Tdh3p or Arc1p. As we mentioned above, daughter budding yeast cells actively monitor their size before passing a commitment step in late G1, called Start, and initiate DNA replication. For any given strain and nutrient environment, a highly reproducible parameter reflecting the timing of Start is the ‘critical size’ (Polymenis, 2022b), defined functionally here as the size at which 50% of the cells are budded. We found that cells lacking either Tdh3p or Arc1p have a larger critical size than otherwise wild-type cells (Figure 5A), consistent with delayed Start. However, the rate at which these cells increased in size was similar to that of wild-type cells (Figure 5B). From asynchronous cultures of these strains, we noticed that cells lacking Tdh3p or Arc1p were bigger (Figure 5C, left panel), and *tdh3*Δ cells also appear to have a larger birth size than their wild-type counterparts (Figure 5C, right panel). Taken together, our results argue for rather specific effects of these mRBPs on size homeostasis (Figure 5C) and cell cycle progression (Figure 5A), which do not arise from severe growth defects (Figure 5B). In particular, loss of Tdh3p delays the G1/S transition, reflected in the larger critical size of *tdh3*Δ cells, and also delays exit from mitosis, accounting for the larger birth size of *tdh3*Δ cells (Figure 5C, right panel).

**Figure 5.**
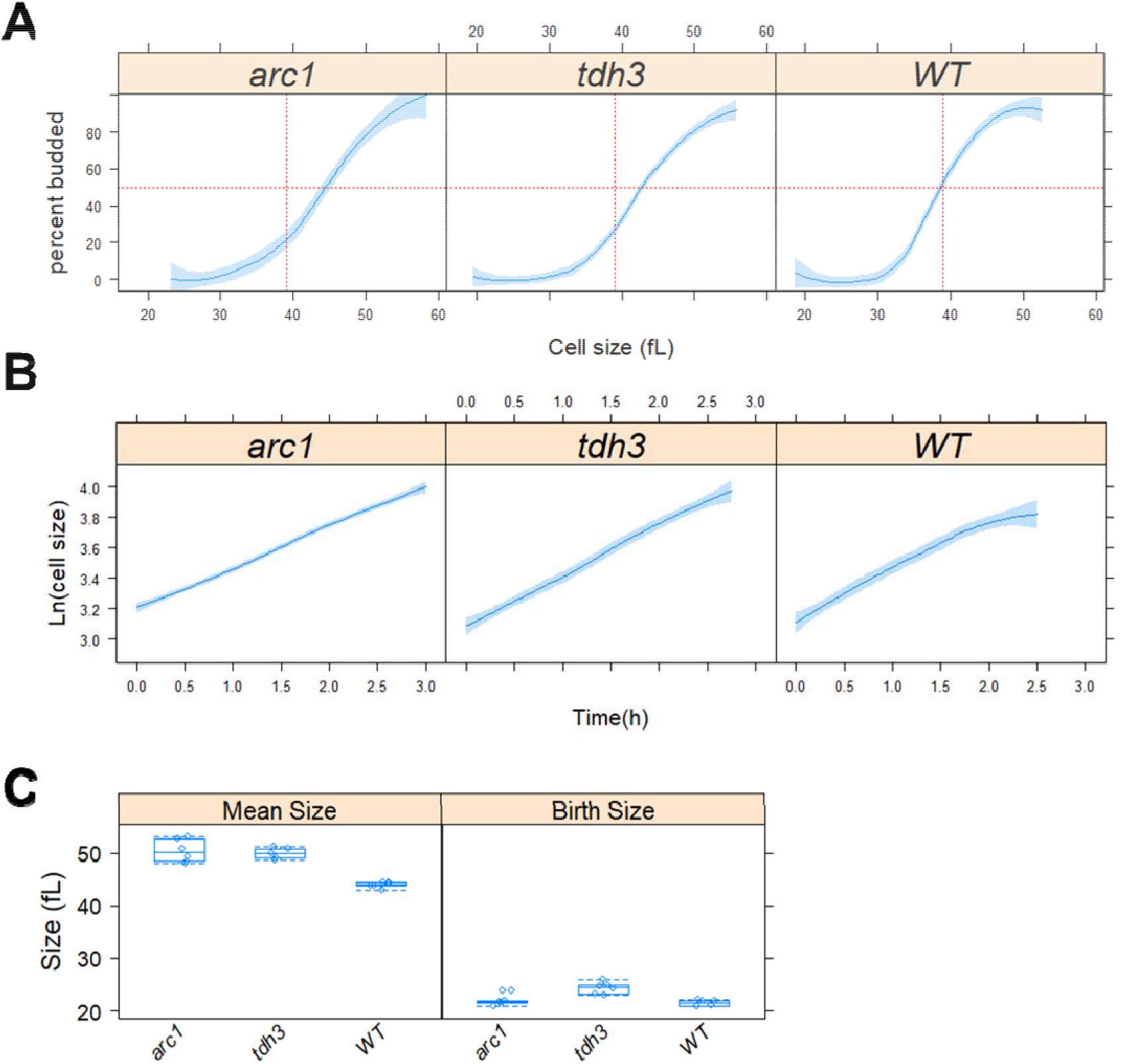
Altered cells size homeostasis and cell cycle kinetics in cells lacking Tdh3p. A, From the synchronous cultures shown in Figure 4B, the percentage of budded cells (y-axis) is shown against the mean cell size (in fL; x-axis). Loess curves and the std errors at a 0.95 level are shown. B, From the same experiments as above, the rate of size increase is indicated from the plots of the Ln-transformed cell size values (y-axis) against time (x-axis). The values used to generate the graphs in A,B are in File S1/Sheet6. C, Box plots showing the mean (left panel) and birth (right panel) size (y-axis) for the indicated strains. Comparisons were made with the nonparametric Kruskal-Wallis rank sum test, and the indicated p-values calculated from the pairwise comparisons using the Wilcoxon rank sum test with continuity correction, using R language functions. The values used to generate the graphs are in File S1/Sheet7.

## Concluding remarks

Our results highlight the role of a key glycolytic enzyme (Tdh3p) through its moonlighting RNA binding properties in the expression of another enzyme (Fas1p) involved in fatty acid biosynthesis. The Tdh3p:*FAS1* interaction is physiologically relevant because it imparts temporal control of Fas1p synthesis, peaking late in the cell cycle. Such interactions could contribute to the temporal compartmentalization of major metabolic pathways during cell division, which based on recent single-cell microscopy studies may be a general feature (Takhaveev *et al*, 2023). Why the Tdh3p:*FAS1* interaction is prominent late in the cell cycle is not clear. Tdh3p is heavily modified by glycosylation (Zielinska *et al*, 2012; Cao *et al*, 2014), ubiquitination (Swaney *et al*, 2013; Back *et al*, 2019), succinylation (Weinert *et al*, 2013; Frankovsky *et al*, 2021), acetylation (Henriksen *et al*, 2012), sumoylation (Bhagwat *et al*, 2021), methylation (Wang *et al*, 2015), and phosphorylation (Lanz *et al*, 2021; Albuquerque *et al*, 2008; MacGilvray *et al*, 2020; Zhou *et al*, 2021; Holt *et al*, 2009; Soulard *et al*, 2010; Rødkær *et al*, 2014). Changes in these posttranslational modifications could modulate the RNA binding properties of Tdh3p. For example, we note that Tdh3p is targeted by the cyclin-dependent kinase (Cdk) at 14 sites (Holt *et al*, 2009). More generally, our results describe the tools and methods to identify cell cycle-dependent interactions between a particular mRNA and proteins in yeast and other systems.

## MATERIALS AND METHODS

A Reagent Table (File S4) is in the Supplementary Files. Where known, the Research Resource Identifiers (RRIDs) are shown in the Reagent Table.

### Media and growth conditions

For bacterial growth during cloning procedures, we used NEB® 5-alpha Competent *E. coli* (High Efficiency) cells from New England Biolabs (Cat#: C2987H), grown in standard LB medium (1% ^w^/_v_ tryptone, 1% ^w^/_v_ NaCl, 0.5% ^w^/_v_ yeast extract, pH 7.0) at 37° with the appropriate antibiotic to maintain plasmid selection. All the *S. cerevisiae* strains used in this study are shown in the Reagent Table. For most experiments, the cells were cultivated in the standard, rich, undefined medium YPD (1% ^w^/_v_ yeast extract, 2% ^w^/_v_ peptone, 2% ^w^/_v_ dextrose), at 30° (Kaiser *et al*, 1994).

### Plasmids and strains

#### dCas13d-dsRBD-APEX2 entry plasmid

The dCas13d-dsRBD-APEX2 plasmid, originally engineered for mammalian expression, was a gift from Alice Ting (Addgene plasmid # 154939; http://n2t.net/addgene:154939; RRID:Addgene_154939), generated as described in (Han *et al*, 2020). With that plasmid as a template, sequences corresponding to positions 1-603, and 604-4071, of the insert were amplified with Phusion® High-Fidelity DNA Polymerase, using primers APEX_1-603_fwd and APEX_1-603_rev, and APEX_604-4071_fwd and APEX_604-4071_rev, respectively. The primers were designed to enable BsmBI/BsaI assembly of the full length (positions 1-4071) dCas13d-dsRBD-APEX2 insert into the entry vector (plasmid YTK001) of the MoClo-YTK plasmid kit (Lee *et al*, 2015), which was a gift from John Dueber (Addgene kit # 1000000061). Note that the insert sequences were amplified in two separate fragments, to remove an internal type IIS restriction site that would interfere with downstream ‘Golden Gate’ cloning strategies. The YTK001 vector and the amplified fragments were subjected to single-pot ‘Golden Gate’ assembly (Engler *et al*, 2008, 2009).

The assembly reaction contained 1 μL of each DNA fragment (from a 20 fmol/μL solution), 1.5 μL T4 ligase buffer (from a 10X solution), 1 μL of T7 ligase, 0.5 μL of restriction enzyme (BsmBI in this case), and water to 15 μL total reaction volume. Unless indicated otherwise, the same reaction composition was used for all assemblies.

The reaction conditions were 30 cycles: 42° for 1 min, 16° for 1 min; followed by 60° for 5 min. The correct assembly of the resulting plasmid (APEX_ENTRY) was verified by sequencing of the entire 4,071 bp insert, with primers FOR_1, FOR_2, FOR_3, FOR_4, FOR_5, FOR_6, REV_1 (see Reagent Table).

#### *FAS1* guide RNA entry plasmids

To design Cas13 RNAs (cRNAs), we used the web-based platform developed by Sanjana and colleagues (Wessels *et al*, 2020). The three RNAs we chose corresponded to positions near the start (positions 277-299), middle (positions 2760-2782), and end (positions 6006-6028) of the *FAS1* mRNA. For each duplex, the two complementary oligonucleotides encoding these sequences (see Reagent Table) were annealed as follows: Each oligonucleotide was resuspended to a final 50μM concentration. Then, 10 μL of each oligonucleotide in the doublex was mixed, to 20 μL total. Annealing was done in the thermocycler, at 95° for 5 min, 55° for 15 min, 25° for 15 min. The annealed oligonucleotides were inserted into the guide RNA entry vector (plasmid YTK050) of the MoClo-YTK plasmid kit (Lee *et al*, 2015), through the ‘Golden Gate’ cloning strategies described in (Akhmetov *et al*, 2018), for 42° for 5 min, 60° for 5 min. The resulting plasmids (FAS1_cRNA-1_ENTRY, FAS1_cRNA-2_ENTRY, FAS1_cRNA-3_ENTRY, respectively), were sequenced with primer t0-ter_FWD (see Reagent Table), to verify the cloning of the RNA guide sequences.

#### Cassette plasmid assembly

The dCas13d-dsRBD-APEX2_001 plasmid was mixed in a single-pot ‘Golden Gate’ assembly with T7 ligase and BsaI, and with plasmids YTK002 (conLS; connector), YTK067 (conR1; connector), YTK009 (*pTDH3*; promoter), YTK063 (*tADH1*; terminator), YTK074 (*URA3*; yeast selection marker), YTK081 (*CEN6/ARS4*; yeast maintenance), YTK083 (AmpR-ColE1; bacterial selection and maintenance), all of which are in the MoClo-YTK plasmid kit (Lee *et al*, 2015), which was a gift from John Dueber (Addgene kit # 1000000061). The assembly reaction conditions were 50 cycles: 37° for 2 min, 16° for 5 min; followed by 60° for 5 min, and 80° for 10 min.

The assembled plasmid (APEX_TU1) encoded a transcriptional unit for dCas13d-dsRBD-APEX2 expression in yeast, from a stably maintained (*CEN6/ARS4*) plasmid. We transformed yeast cells (BY4742 strain) with the APEX_TU1 plasmid. The ends of the insert in the plasmid were sequenced with primers AmpR-FWD and pBR322ori-FWD (see Reagent Table). Then, we verified that the transformants express a protein recognized by an anti-V5 antibody conjugated with HRP (Invitrogen Cat #R96125; see Reagent Table; used at an 1:5,000 dilution) with an apparent molecular mass of 153,650.71 Da, expected for the dCas13d-dsRBD-APEX2 protein (not shown).

The *FAS1* guide RNA cassettes were assembled individually, as described above for APEX_TU1, with T7 ligase and BsaI. Each reaction contained the FAS1_cRNA entry plasmid of interest and plasmids YTK003 (conL1; connector), YTK072 (conRE; connector), YTK083 (AmpR-ColE1; bacterial selection and maintenance), from the MoClo-YTK plasmid kit, yielding plasmids FAS1-1_TU2, and FAS1-2_TU2, respectively. The correct inserts were validated by sequencing, with primers AmpR-FWD and pBR322ori-FWD (see Reagent Table).

To generate FAS1-3_TU2, we first PCR-amplified the corresponding insert using FAS1-3_ENTRY as a template, and primers 050_L1 and 050_RE (see Reagent Table; which encode the appropriate BsmBI sites for the next bicistronic assembly).

#### Bicistronic plasmid assembly and yeast expression

To drive expression in yeast of dCas13d-dsRBD-APEX2 and each of the *FAS1* cRNAs off the same plasmid, we combined with T7 ligase and BsmBI plasmids APEX_TU1, YTK096, and one of FAS1-1_TU2 plasmid, FAS1-2_TU2 plasmid; or the FAS1-3_TU2 PCR fragment. These assembly reaction conditions were 50 cycles: 42° for 2 min, 16° for 5 min; followed by 60° for 5 min, and 80° for 10 min. They yielded plasmids APEX-FAS1-1_INT, APEX-FAS1-2_INT, APEX-FAS1-3_INT; respectively. Correct assembly was validated by sequencing, with primers FOR_6 and 050_RE (see Reagent Table). Plasmids APEX-FAS1-1_INT, APEX-FAS1-2_INT, APEX-FAS1-3_INT were each digested with NotI and used to transform strain BY4742, yielding strains SCMSP244, SCMSP245, and SCMSP246, respectively. Each of these strains carry an integration of the bicistronic assembly into the *URA3* locus. The strains were validated by APEX protein expression, through immunoblotting against the V5 epitope, and sequencing of the chromosomal locus for the presence of the correct gRNA. Lastly, we also ensured that the APEX fusion was active in these strains, using the Amplex^Ⓡ^ Red assay described previously (Turnšek *et al*, 2021), generating the strongly fluorescent resorufin (Dębski *et al*, 2016), as shown in Figure 1.

#### Yeast mutants

Single gene haploid deletion strains, lacking *ARC1*, *SRO9*, *SSD1*, or *TDH3*, were commercially available (see Reagent Table). Their genotype was validated by PCR, to confirm that the gene of interest was absent and replaced by the appropriate marker. These strains were crossed with a commercially available *FAS1-TAP* strain (see Reagent Table), sporulated and dissected to obtain the corresponding haploid deletion mutant of the mRNA binding protein carrying a *FAS1-TAP* allele.

### Centrifugal elutriation and cell size measurements

All methods have been described previously (Hoose *et al*, 2012; Soma *et al*, 2014; Blank *et al*, 2020, 2017b; Maitra *et al*, 2020). Briefly, to collect enough cells for the LC-MS/MS measurements after proximity labeling, elutriated G1 cells were allowed to progress in the cell cycle until they reached the desired cell size. At that point, they were quenched (with 100 µg/ml cycloheximide) and frozen away, and later pooled with cells of similar size. Overall, we had to collect 54 individual samples, to generate the 18 pools shown in Figure 3B.

For other elutriation experiments (e.g., see Figures 4,5), only an early G1 elutriated fraction was collected, from which samples were taken at regular intervals as the cells progressed in the cell cycle.

### Proximity labeling reactions

For each labeling reaction, 1E+09 cells collected and stored at −80° in freezing buffer (15% glycerol, 150 mM potassium acetate, 2 mM magnesium acetate, 20 mM HEPES/sodium hydroxide pH 7.2, 0.5% (^w^/_v_) glucose, 100 μg/mL cycloheximide) were thawed on ice. The cells were washed in 10 mL 0.1M MES/sodium hydroxide pH 6.5, 100 μg/mL cycloheximide, resuspended in 2 mL of this buffer containing 0.01% digitonin (10 μL added from a 20 mg/mL digitonin stock in DMSO), and incubated in a 30° shaking water bath for 30 min (Cordeiro & Freire, 1995). The cells were then collected by a brief centrifugation, washed with 10 mL of ice-cold 1.2M sorbitol/PBS solution, resuspended in 2 mL of 1.2M sorbitol/PBS containing 2.5 mM phenol-biotin (10 μL added from a 0.5M stock in DMSO, stored at −80°), 20 μL from a 20 U/μL stock of SUPERase•In RNase Inhibitor, 20 μL from a 10 mg/mL stock of cycloheximide, and protease inhibitors, and incubated on ice for 90 min (Turnšek *et al*, 2021). About 15 min before the end of the 90 min incubation on ice, a quenching solution was prepared by mixing the following: 200 μL Trolox (from a 0.5M stock in DMSO, stored at −80°); 200 μL sodium azide (from a 1M stock in water, stored at −80°); 2 mL of a 10 mM sodium ascorbate solution in PBS, prepared fresh. Also before the end of the 90 min incubation on ice, a 0.2 M stock of hydrogen peroxide was prepared (by dilution of a 30% (9.8 M) hydrogen peroxide solution; stored at 4°). At the end of the 90 min incubation on ice, the cells were exposed to 2mM hydrogen peroxide (20 μL were added to the 2 mL cell suspension from the 0.2 M solution), vortexed briefly, and incubated on ice for 2 min. The reaction was stopped by adding 2 mL of the freshly prepared quenching solution described above. The cells were collected by centrifugation, washed with 10 mL of TBS, pH 7.5, and resuspended in 3 mL of TBS, pH 7.5. Then 1.5 mL of glass beads were added to each tube, to break the cells with 6 cycles of 30 sec vortexing-30 sec on ice. The cells were centrifuged for 10 min in the cold, and the supernatants transferred to 15 mL screw-cap tubes.

To isolate the biotinylated proteins for mass spectrometry, 0.2 mL of beads (Dynabeads™ MyOne™ Streptavidin C1; ThermoFisher, Cat#: 65001) were added to each tube, and Incubated on a rotisserie mixer for 1 h at room temperature. A magnetic rack was used to isolate the beads and remove the supernatant. The beads were washed twice with 10 mL TBS, 2 M urea, pH 7.5, once with 10 ml 0.1 M ammonium carbonate pH 7.7, and resuspended in 0.5 ml 0.1 M ammonium carbonate pH 7.7.

The APEX labeling reactions shown in Figure 2 were done as described above but from 1E+08 cells, with all the reaction volumes reduced 10-fold, and then processed for immunoblotting as described below.

### LC-MS/MS

The beads were washed three times with 200 μL of 25 mM ammonium bicarbonate. After the final wash, 200 μL of 25 mM ammonium bicarbonate was added along with 2 μg of proteomics-grade trypsin (2 μL of a 1 μg/μL solution) and incubated for a day at 37° with intermittent vortexing. An aliquot of the supernatants from the resultant samples were diluted two-fold in Solvent A (95/5% water/acetonitrile containing 0.1% formic acid); and 1 μL injected for analysis by LC-MS/MS. The nanoLC-MS consisted of an UltiMate 3000 Nano LC System and an LTQ-Orbitrap Elite mass spectrometer (Thermo Fisher, San Jose, CA). Reversed-phase liquid chromatography was performed using a homemade 33 cm × 75 μm ID column packed with XBridgeTM BEH C18 media (2.5 μm, 130 Å). The flow rate was maintained at 200 nL/min. Solvent A and B (95/5% acetonitrile/water containing 0.1% formic acid) were used to establish the 160 min gradient elution timetable: isocratic at 5% B for 30 min, 5-55% B over 70 min, followed by 55-99% B in 5 min where it was maintained for 10 min, and finally returned to 5% B over 5 min for a 40 min re-equilibration time. The LTQ-Orbitrap Elite mass spectrometer instrument was operated in positive mode with a 2.6 kV applied spray voltage. The temperature of the ion transfer capillary was 300°. One microscan was set for each MS and MS/MS scan. A full scan MS acquired in the range 300 ≤ m/z ≤ 2000 was followed by 10 data dependent MS/MS events on the 10 most intense ions. The mass resolution was set at 60000 for full MS. The dynamic exclusion function was set as follows: repeat count, 1; repeat duration, 30 s; exclusion duration, 30s. HCD was performed using normalized collision energy of 35% and the activation time was set as 0.1 ms. Mascot software (Matrix Science, Boston, MA) was used for protein identification and quantitation. The mass spectrometry proteomics data have been deposited to the ProteomeXchange Consortium via the PRIDE (Perez-Riverol *et al*, 2022) partner repository with the dataset identifier PXD041908. To identify proteins that were preferentially associated with the *FAS1* mRNA in the G1 phase or in the G2 phase, each strain was analyzed separately with the robust bootstrap ANOVA. The input values used in each case are shown in File S2. The output values and the fold-change are in File S1/Sheet3, and plotted in the volcano plots shown in Figure 3C.

### Immunoprecipitations

#### dCas13d-dsRBD-APEX2-V5

Exponentially proliferating cells were quenched with 100 μg/mL cycloheximide. They were then collected by centrifugation and washed with freezing buffer (15% glycerol, 150 mM potassium acetate, 2 mM magnesium acetate, 20 mM HEPES/sodium hydroxide pH 7.2, 0.5% (^w^/_v_) glucose, 100 μg/mL cycloheximide), resuspended in the same freezing buffer (1E+08 cells in 60 μL buffer), and stored at −80° until further use (Singer-Krüger *et al*, 2020). The cells were washed in 1 mL of RIP buffer (150 mM KCl, 25 mM Tris pH 7.4, 5 mM EDTA, 0.5 mM DTT, 0.5% IGEPAL® CA-630), and resuspended in 0.6 mL of RIP buffer containing 100 U/mL RNAse inhibitor SUPERase•in™ (ThermoFisher; Cat#: AM2694, see Reagent Table) and protease inhibitor cocktail (Millipore Sigma; Cat#: 11836170001) (the RNAse and protease inhibitors were added fresh). About 0.250 mL of glass beads were added and vortexed at the maximum speed for 30 sec, then placed on ice for 30 sec. The vortex-ice cycle was repeated for a total of six times, to break the cells. The supernatant was collected after a centrifugation at 5,000 rpm for 5 min, and clarified with another centrifugation at 12,000 rpm for 2 min. The clarified supernatant was removed and 0.15 mL was stored at −80°, to serve as ‘input’ control. To the rest, 10 μL of agarose-α-V5 beads were added, and incubated at 4° on a tube rotator for 1-2 h. The beads were pelleted at 1,000 rpm for 1 min, and the supernatant was removed. The beads were washed with 0.5 mL RIP buffer, and pelleted as before. Two additional such washes were performed, and the beads were resuspended in 130 μL of RIP buffer and stored at −80°, prior to RNA isolation and ddPCR.

#### TAP-tagged proteins

Because many RNA binding proteins are found in stress granules, we adapted an approach described previously to generate cell extracts that recover such structures (Jain *et al*, 2016). Briefly, for each TAP-tagged strain, cells from 1 L of culture (in YPD) was harvested and resuspended in 10 mL of lysis buffer (10mM Tris-HCl pH 7.5, 100mM sodium chloride, 1.5mM magnesium chloride, 0.5% NP-40), with 1:5,000 Antifoam emulsion and protease inhibitor cocktail added. 5 mL of glass beads were added and the cells were lysed by 3 cycles of vortexing for 2 min followed by 2 min on ice. The lysates were centrifuged at 850 *g* for 2 min and the supernatants collected. Then 0.2 mL of washed IgG Sepharose 6 Fast Flow beads (Millipore Sigma, Cat#: GE17-0969-01) were added to each sample and incubated on a rotisserie mixer for 0.5 h at room temperature. The beads were washed 3 times with a buffer containing 10mM Tris-HCl pH 7.5, 100mM sodium chloride, 1.5mM magnesium chloride, 0.1% NP-40. The beads were resuspended in 500 μL of the same buffer and stored at −80°, prior to RNA isolation and ddPCR.

### Immunoblotting

For the samples shown in Figure 2, the cells were collected by centrifugation, resuspended in 0.1 mL 0.1N sodium hydroxide, and incubated at room temperature for 5 min. An equal volume of 2x Laemmli buffer (65.8 mM Tris-HCl, pH 6.8, 2.1% SDS, 26.3% ^w^/_v_ glycerol, 0.01% bromophenol blue) was added to the samples prior to SDS-PAGE. Biotinylated proteins were detected with a streptavidin-HRP conjugate (Millipore-Sigma, Cat#: OR03L-200UG), used at 1:2,000 dilution (see Reagent Table).

To detect Fas1p-TAP (see Figure 4), protein extracts were made as described previously (Amberg *et al*, 2006), and resolved on 6% Tris-Glycine SDS-PAGE gels. To detect Fas1p-TAP with the PAP reagent (Millipore-Sigma; Cat#: P1291, used at 1:5,000 dilution), we used immunoblots from extracts of the indicated strains, as we described previously (Blank *et al*, 2017b, 2020; Maitra *et al*, 2020, 2022). Loading was measured with an anti-Pgk1p primary antibody (ThermoFisher; Cat#: 459250) used at 1:5,000 dilution, and a secondary anti-mouse HRP-conjugated antibody (Abcam; Cat#: ab205719) used at 1:5,000 dilution. Imaging and quantification was done as described previously (Blank *et al*, 2017b, 2020; Maitra *et al*, 2020, 2022).

### Digital droplet PCR (ddPCR)

All methods have been described previously (Maitra *et al*, 2022).

### Statistical analysis, sample-size and replicates

For sample-size estimation, no explicit power analysis was used. There was also no randomization or blinding during sample analysis. All the replicates in every experiment shown were biological ones, from independent cultures. A minimum of three biological replicates was analyzed in each case, as indicated in each corresponding figure’s legends. The robust bootstrap ANOVA was used to compare different populations via the t1waybt function, and the posthoc tests via the mcppb20 function, of the WRS2 R language package (Wilcox, 2011; Mair & Wilcox, 2020). Note that with the robust bootstrap ANOVA exact p values <0.0001 were not calculated. We also used non-parametric statistical methods, as indicated in each case. The Kruskal-Wallis and posthoc Nemenyi tests were done with the posthoc.kruskal.nemenyi.test function of the PMCMR R language package. No data or outliers were excluded from any analysis.

## Data availability

Strains and plasmids are available upon request. The authors affirm that all data necessary for confirming the conclusions of the article are present within the article, figures, and tables. LC-MS/MS data are available via ProteomeXchange with identifier PXD041908. The Supplementary Material includes the following:

File S1: Source data for the graphs in all Figures.

File S2: Source input data for the ANOVA analysis of the LC-MS/MS data.

File S3: Source immunoblots for Figure 4.

File S4: Reagent Table

## ACKNOWLEDGEMENTS

This research was supported by a grant from the National Institutes of Health (GM123139) to M.P.. LC-MS/MS analyses were carried out at the UTSA Mass Spectrometry & Proteomics Core.

## AUTHOR CONTRIBUTIONS

M.P. conceived and designed the study. W.P.G. performed the LC-MS/MS experiments. H.M.B. and M.P. performed all other experiments. All authors evaluated and discussed the data. M.P. wrote the first draft of the manuscript, which was then edited by all authors.

## CONFLICT OF INTEREST

The authors have no conflicts of interest to declare that could be perceived to influence the presentation or interpretation of the data.

## Notes

### Competing Interest Statement

The authors have declared no competing interest.

