## Supplementary material for "Targeting APEX2 to the mRNA encoding fatty acid synthase β in yeast identifies proteins that bind and control its translational efficiency in the cell cycle": File S3

**File S3. Source immunoblots for Figure 4.**

To the right of each blot is the background subtracted signal captured by the imaging system, and used to quantify the band intensities. For the wild-type cells, in addition to the three experiments shown, we included in the analysis immunoblots we had published previously (Blank et al, 2017b; Maitra et al, 2022).

Strain: WT (FAS1-TAP)

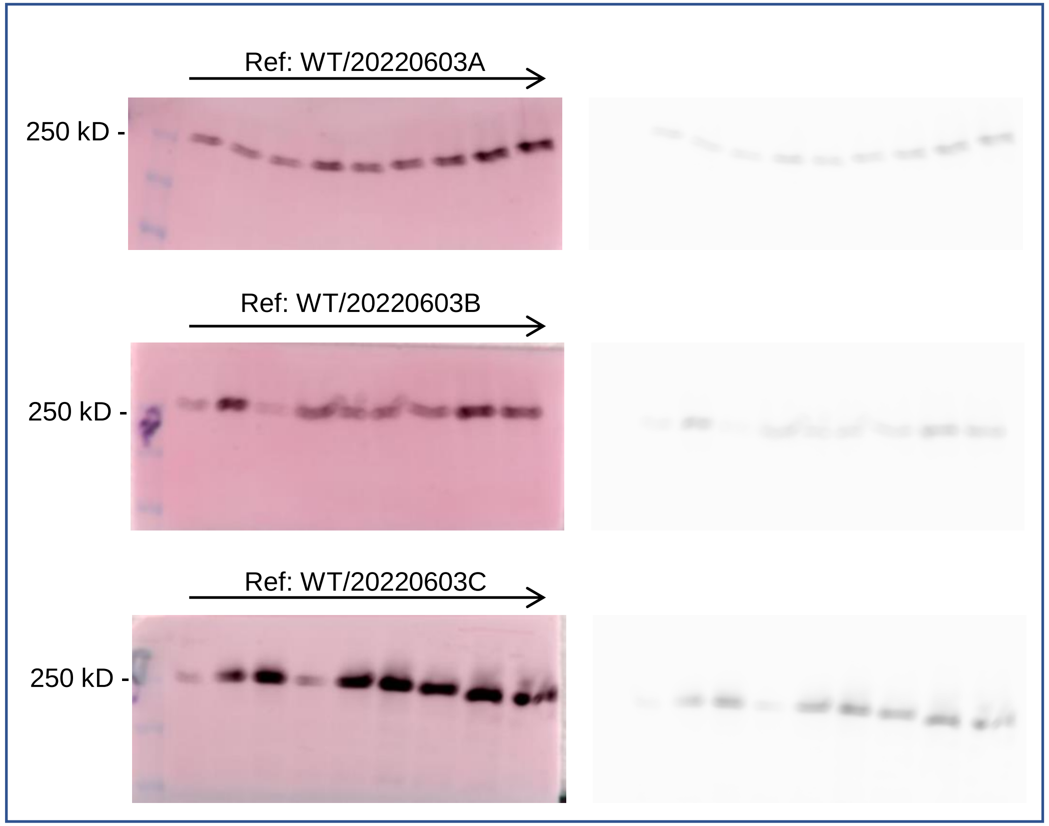


α-Fas1-TAP

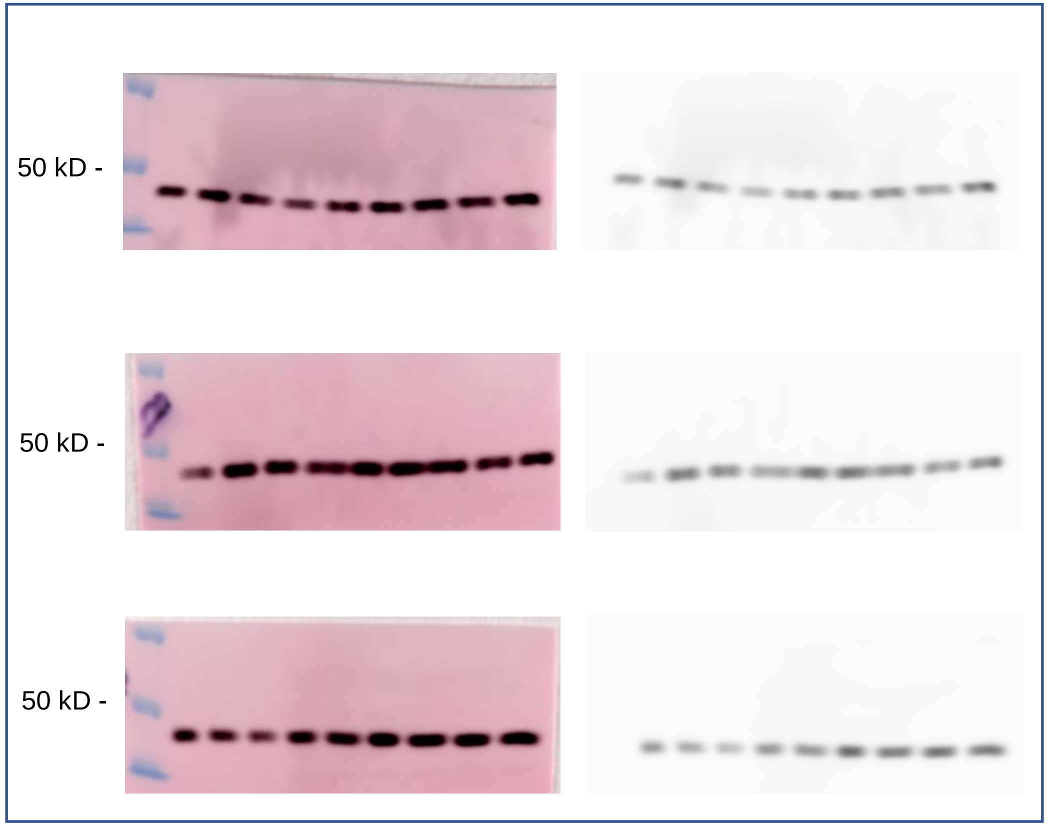


α-Pgk1p

Strain: tdh3 (FAS1-TAP)

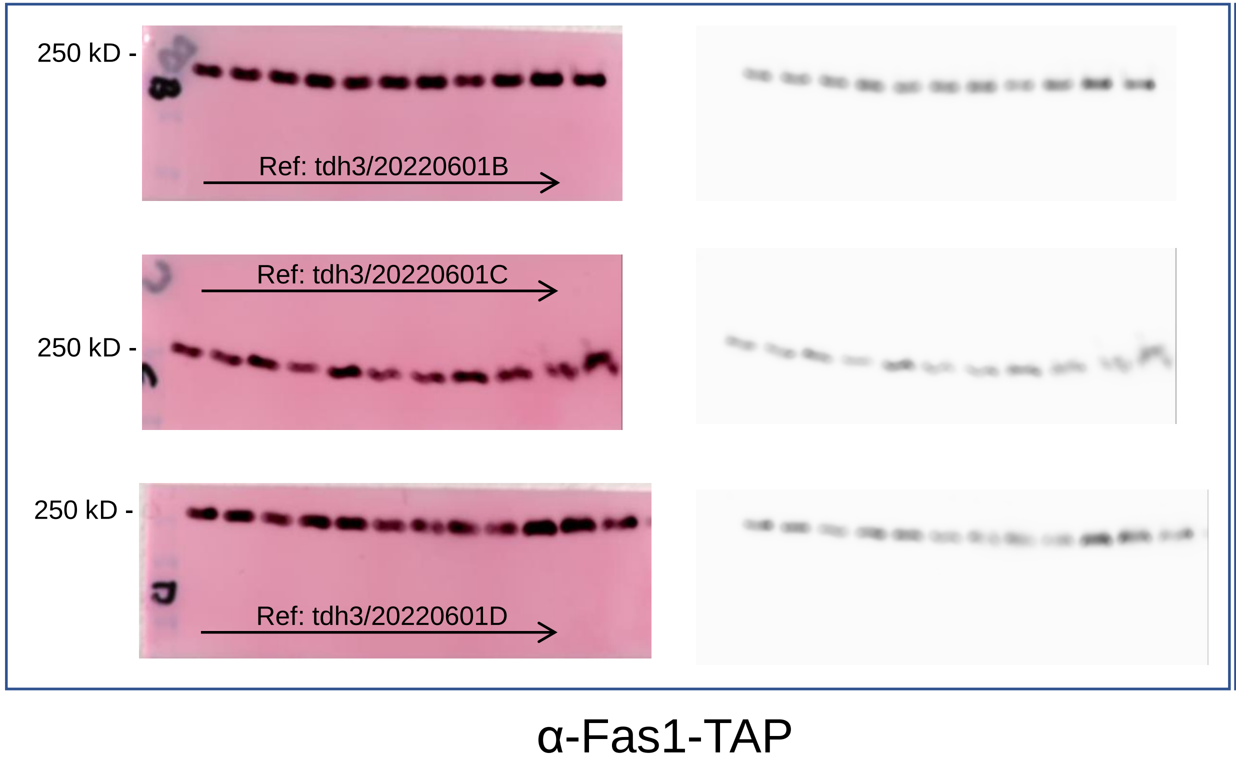


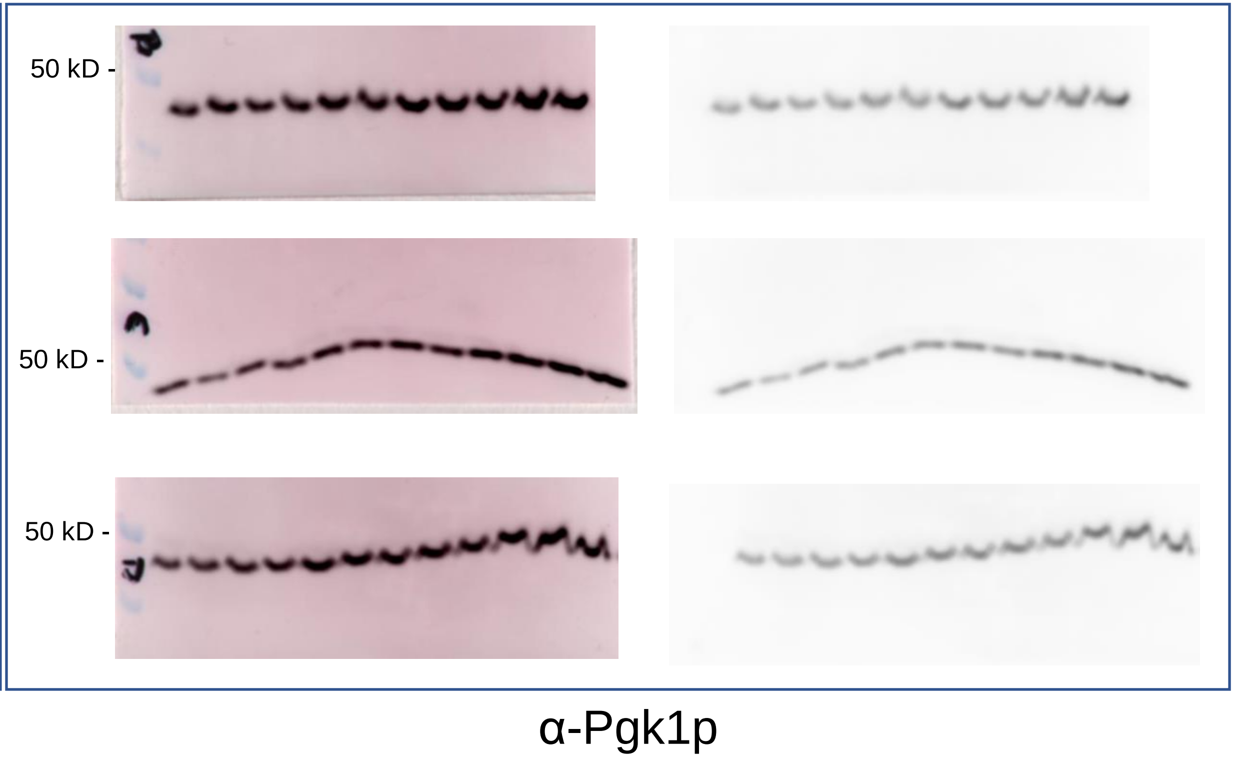

### Strain: arc1 (FAS1-TAP)

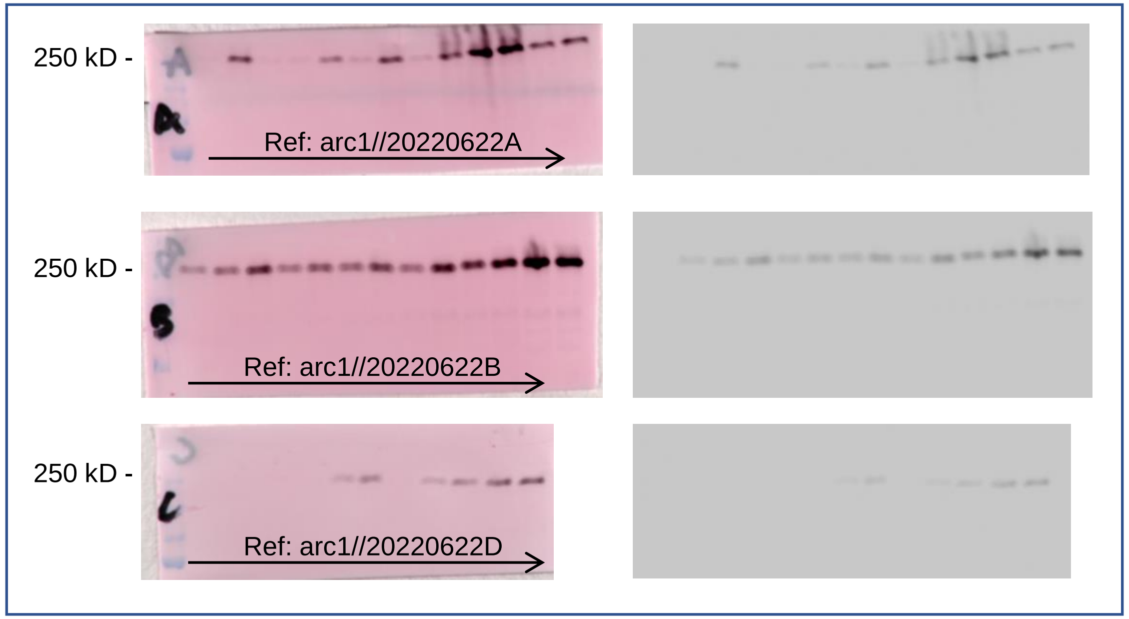


α-Fas1-TAP

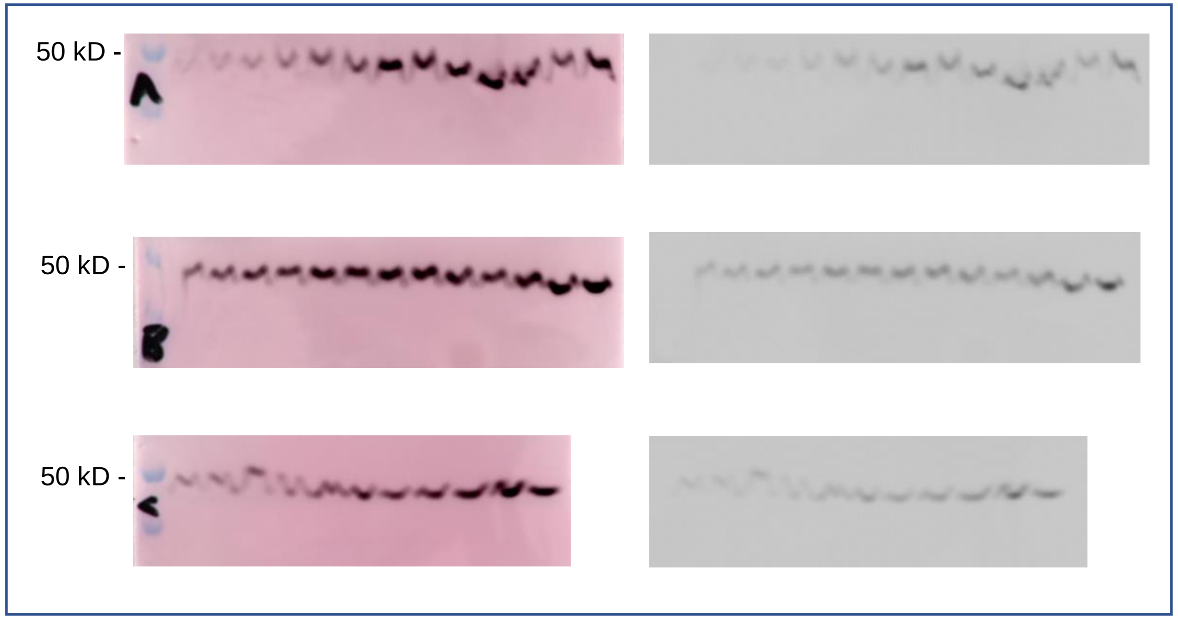


α-Pgk1p
